# Population genomics of a thermophilic cyanobacterium revealed divergence at subspecies level and possible adaptation genes

**DOI:** 10.1101/2024.08.22.609105

**Authors:** Hsin-Ying Chang, Hsi-Ching Yen, Hsiu-An Chu, Chih-Horng Kuo

## Abstract

**Background:** Cyanobacteria are diverse phototrophic microbes with ecological importance and potential for biotechnology applications. One species of thermophilic cyanobacteria, *Thermosynechococcus taiwanensis*, has been studied for biomass pyrolysis, estrogen degradation, and the production of bioethanol, monosaccharide, and phycocyanin. To better understand the diversity and evolution of this species, we sampled across different regions in Taiwan for strain isolation and genomic analysis.

**Results:** A total of 27 novel strains were isolated from nine of the 12 hot springs sampled and subjected to whole genome sequencing. Including strains studied previously, our genomic analyses encompassed 32 strains from 11 hot springs. Genome sizes among these strains ranged from 2.64 to 2.70 Mb, with an average of 2.66 Mb. Annotation revealed between 2,465 and 2,576 protein-coding genes per genome, averaging 2,537 genes. Core-genome phylogeny, gene flow estimates, and overall gene content divergence consistently supported the within-species divergence into two major populations. While isolation by distance partially explained the within-population divergence, the factors driving divergence between populations remain unclear. Nevertheless, this species likely has a closed pan-genome comprising approximately 3,030 genes, with our sampling providing sufficient coverage of its genomic diversity. To investigate the divergence and potential adaptations, we identified genomic regions with significantly lower nucleotide diversity, indicating loci that may have undergone selective sweeps within each population. We identified 149 and 289 genes within these regions in populations A and B, respectively. Only 16 genes were common to both populations, suggesting that selective sweeps primarily targeted different genes in the two populations. Key genes related to functions such as carbon fixation, photosynthesis, motility, and ion transport were highlighted.

**Conclusions:** This work provides a population genomics perspective on a hot spring cyanobacterial species in Taiwan. Beyond advancing our understanding of microbial genomics and evolution, the strains collected and genome sequences generated in this work provide valuable materials for future development and utilization of biological resources.

## Background

Cyanobacteria are a diverse group of photosynthetic microorganisms that play crucial roles as primary producers and are vital components of aquatic ecosystems (Sánchez-Baracaldo et al. 2022). In addition to their ecological significance, these organisms can be harnessed for bioremediation (Touliabah et al. 2022), biofuel production (Singh et al. 2023), and the synthesis of high-value secondary metabolites (Kultschar et al. 2018). Thermophilic cyanobacteria, in particular, are highly valued in industrial applications due to their production of thermostable enzymes and proteins (Patel et al. 2019).

One genus of thermophilic cyanobacteria, *Thermosynechococcus*, thrive in non-acidic hot springs with a temperature range of about 50-65°C across several Asian countries and have attracted much research attention (Everroad et al. 2012; Nishida et al. 2018; Tang et al. 2018; Ward et al. 2019; Liang et al. 2019; Prondzinsky et al. 2021). A recent update of *Thermosynechococcus* taxonomy identified eight species within this genus (Tang et al. 2024). One species, tentatively named as *Thermosynechococcus taiwanensis*, is of particular interest. Two strains belonging to this species, CL-1 and TA-1, were shown to have potential in biomass pyrolysis (Hsueh et al. 2007), bioethanol production (Su et al. 2013), monosaccharide production and estrogen degradation (Chang et al. 2021), and phycocyanin production (Leu et al. 2013; Narindri Rara Winayu et al. 2022). Our genomic characterizations of these two strains revealed their genetic divergence at within- and between-species levels, such as genes involved in transportation, nitric oxide protection, urea utilization, kanamycin resistance, restriction modification system, and chemotaxis (Cheng et al. 2020, 2022). Phylogenetically, *T. taiwanensis* is most closely related to two metagenome-assembled genomes from India, which represent a novel species that has yet to be cultivated (Cheng et al. 2022; Tang et al. 2024). Among the cultivated members of *Thermosynechococcus, T. taiwanensis* is most closely related to strain E542 from China. The strain E542 was initially identified as *Thermosynechococcus elongatus* (Liang et al. 2019) and recently reclassified as *Thermosynechococcus* sichuanensis (Tang et al. 2024).

To further improve our understanding of *T. taiwanensis*, we expanded the sampling to 12 additioanl hot springs distributed across Taiwan and isolated 27 novel strains in this study. Through whole-genome characterization of these newly isolated strains and comparative analysis with other strains, the population genomics work reported here greatly improved the characterization of their genetic diversity and allowed for the inference of genes that likely underwent selective sweeps. In addition to the contribution to microbial genomics and evolution, the collection of novel bioresources and knowledge regarding their genetic diverstiy can provide a strong foundation for future developments of biotechnology applications.

## Methods

### Biological samples and growth condition

Field samples were collected from 12 hot springs in Taiwan during the period of from June 2022 to April 2023 (**Figure 1 and Table S1**). At each site, the water temperature and pH were recorded, and the green biofilm on rocks under water was scraped into sterilized 1.5 mL polypropylene centrifuge tubes. After being transported to the laboratory, the biological materials were maintained in BG-11 liquid medium (Rippka et al. 1979) supplemented with 20 mM TES (pH 8.0) under continuous white LED light (20 μmol photons m^−2^s^−1^) at 45°C. For strain isolation, the culture was streaked out on BG-11 solid agar plates (5 mM TES, pH 8.0) containing 12.5 μg/ml of kanamycin and grown under the same lighting and temperature. Single colonies were identified and streaked out for at least three more times to obtain pure strains. The assignment of these strains to the genus *Thermosynechococcus* was confirmed by colony PCR and Sanger sequencing of their 16S rRNA gene using the primer pair 27F (5’-AGAGTTTGATCMTGGCTCAG-3’) and 1492R (5’-GGTTACCTTGTTACGACTT-3’).

**Figure 1.**
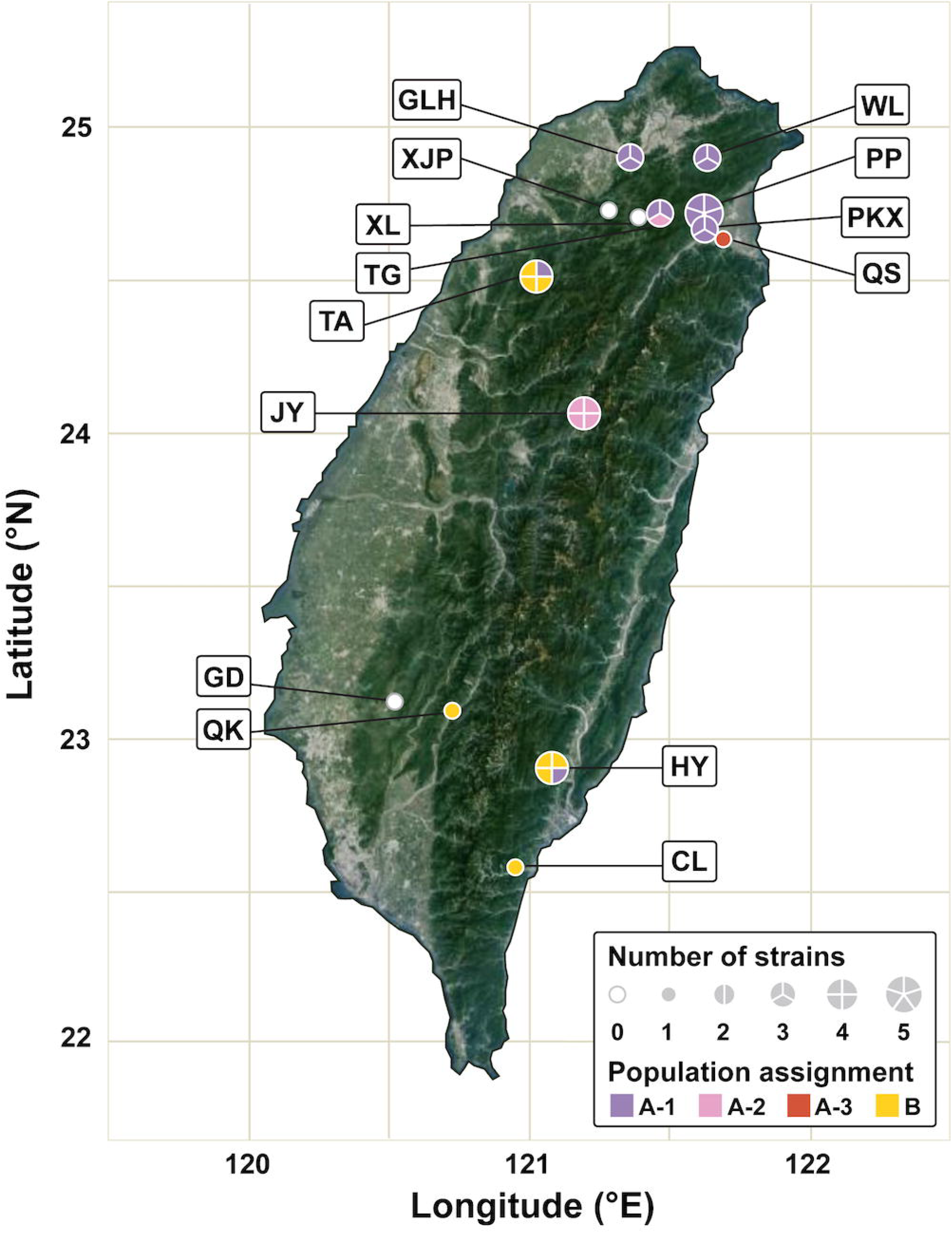
The hot springs in Taiwan sampled in this study. For each site, a pie chart is used to indicate the number of strains characterized and the population assignment of individual strains based on 99% average nucleotide identity (ANI). Abbreviations of the hot spring names: CL, Chinlun; GD, Gueidan; GLH, Galahe; HY, Hongye; JY, Jingying; PKX, Paikuxi; PP, Pengpeng; QK, Qikeng; QS, Quishui; TA, Taian; TG, Taigang; WL, Wulai; XJP, Xiaojinping; XL, Xiuluan.

### Genome sequencing

The procedures for whole-genome shotgun sequencing and analysis were based on those described in our previous studies (Cheng et al. 2020, 2022). More detailed information is provided in the following paragraphs. Unless stated otherwise, the methods were based on the cited references, the kits were used according to the manufacturer’s instructions, and the bioinformatics tools were used with the default settings.

For the extraction of genomic DNA, each strain was grown in 150 ml of BG-11 liquid medium supplemented with 20 mM TES (pH 8.0) and 100 mM sodium bicarbonate at 45°C for 4-5 days. For each sample, approximately 1 g of cells were collected by centrifugation (4,000 x g for 5 min), ground in liquid nitrogen, then mixed with 5 ml of extraction buffer (0.9% Sodium bisulfite, 0.35 M Sorbitol, 0.1 M Tris-base, and 5 mM EDTA, pH 8.0), 5 ml of nuclei lysis buffer (2% CTAB, 2 M NaCl, 200 mM Tris-base, and 50 mM EDTA, pH 7.5), 2 ml of 5% Sarkosyl solutionthen, and incubated at 65°C for cell lysis. After incubation for 30 min, 0.8 ml of the solution was mixed well with an equal volume of chloroform:isoamyl alcohol (24:1). Following centrifugation at 10,000 x g for 1 min, the mixture was separated into two phase layers. The water phase layer was transferred to a clean sterilized 1.5 ml tube and added with 0.6 volume of isopropanol for DNA precipitation, then centrifugated at 10,000 x g for 10 min to collect the DNA pellet. The DNA pellet was washed with 0.8 ml 75% ethanol twice and air dried at room temperature. The pellet was then dissolved in 0.05 ml deionized water and processed a further purification with the DNeasy Blood and Tissue Kit (No. 69504; Qiagen, Germany). The purified genomic DNA samples were assessed by using the NanoDrop 2000 spectrophotometer (ThermoFisher, United States) and 1% agarose gel electrophoresis for quantification and quality control. Shearing and size selection was not performed for the DNA samples.

For whole-genome shotgun sequencing, the Illumina platform was used for all strains and the Oxford Nanopore Technologies (ONT) platform was used for three representative strains (i.e., HY596, JY1339, and PP45). For Illumina sequencing, the paired-end libraries was prepared by using KAPA LTP Library Preparation Kit (KK8232; Roche, Switzerland) without amplification, then sequenced by using a NovaSeq 6000 sequencer. The sequencing results were processed using Trimmomatic v0.39 (Bolger et al. 2014) to remove adapter sequences and low-quality reads. For ONT sequencing, the library was prepared by using the ONT Ligation Kit (SQK-LSK109) and sequenced by using MinION (FLO-MIN106; R9.4 chemistry and MinKNOW Core v3.6.0). Guppy v6.5.7 was used for basecalling with the super accuracy mode.

For the three representative strains with both Illumina and ONT reads, the *de novo* assembly was conducted by using Unicycler v0.4.9b (Wick et al. 2017). For the remaining strains with only Illumina reads, the de novo assembly was conducted by using Velvet v1.2.10 (Zerbino and Birney 2008). For validation, the Illumina and ONT raw reads were mapped to the resulting assemblies using BWA v0.7.17 (Li and Durbin 2009) and Minimap2 v2.15 (Li 2018), respectively. The mapping results were programmatically checked using SAMtools v1.2 (Li et al. 2009) and manually inspected using IGV v2.16.2 (Robinson et al. 2011). The finalized assemblies were submitted to the National Center for Biotechnology Information (NCBI) and annotated with their Prokaryotic Genome Annotation Pipeline (PGAP) v6.5 (Tatusova et al. 2016).

### Comparative genomics

For comparative analysis, the complete genome assemblies of five additional strains isolated from Taiwan were obtained from GenBank (Sayers et al. 2022). Furthermore, strain E542 from China was included as as the outgroup. The accession numbers and other relevent information of all 33 genome assemblies included in this study are provided in **Table S2**.

For assessment of overall genome similarity, all possible pairwise comparisons were conducted using FastANI v1.1 (Jain et al. 2018) to calculate the percentage of genome fragments mapped and the average nucleotide identity (ANI). For comparison of gene content among all strains, the homologous gene clusters (HGCs) among all strains were identified by using OrthoMCL v1.3 (Li et al. 2003). For principal coordinates analysis of gene content dissimilarity, the matrix of HGC counts was coverted into a Jaccard distance matrix using the VEGAN package v2.6-4 (Oksanen et al. 2022) in R v4.2.3 (R Core Team 2019), then processed using the PCOA function in the APE package v 5.7-1 (Popescu et al. 2012). For phylogenetic analysis, MUSCLE v3.8.31 (Edgar 2004) was used for multiple sequence alignments and PhyML v3.3 (Guindon et al. 2010) was used for maximum-likelihood inference.

For population delineation, PopCOGenT (Arevalo et al. 2019) was used to infer a gene flow network and to conduct clustering of strains. After the populations were defined, genomic regions exhibiting within-population nucleotide diversity levels below the 95% confidence interval were inferred as loci that have experienced selective sweeps. Genes located at these loci were extracted and further examined by using the COG classifier and the KEGG BRITE database (Kanehisa et al. 2010) to investigate their functions.

The pan-genome analysis was conducted by using Roary v3.13.0 (Page et al. 2015) with the cutoff value for sequence identity at the BLASTP (Boratyn et al., 2013) step set to 90%. The inference of pan-genome size was conducted based on Heap’s law (Tettelin et al. 2008), represented by the formula: n = κN^γ^, with n being the pan-genome size, N being the number of genomes included in the analysis, and κ and γ being fitting parameters. A total of 1,000 times of sub-sampling were conducted to estimate the parameters.

The R package ggplot2 v3.4.4 (Wickham 2016) was used for data visualization and the R package stats v4.3.3 was used for linear regression analysis.

## Results and discussion

### Genomic diversity

From the 12 hot springs sampled in this study, 27 new strains were isolated from nine sites (**Figure 1 and Table S1**). Among these, we obtained complete genome assemblies for 22 strains. For the remaining five strains, high-quality draft assemblies with contig N50 ranging from 158.3 to 420.8 kb were obtained. Combined with five additional complete assemblies from GenBank, a total of 32 strains isolated from 11 different hot springs in Taiwan were included in this population genomics study (**Table S2**). The genome sizes of these strains range from 2.64 to 2.70 Mb, with an average of 2.66 Mb. Notably, strain PP45 harbors a 2.3 kb circular plasmid, whereas no plasmids were detected in the other strains. Based on the annotation, the number of intact protein-coding genes ranges from 2,465 to 2,576 among these genomes, with an average of 2,537 per genome.

Based on the commonly accepted criterion of 95% ANI as the species boundary (Jain et al. 2018; Konstantinidis 2023), all 32 strains collected in Taiwan were assigned to the same species-level taxon, namely *T. taiwanensis* (**Figure 2A**). Compared to the most closely related *Thermosynechococcus* lineage that has been cultivated, namely *T. sichuanensis* E542 from China, the between-species ANI values ranged from 91% to 92%. This finding further confirmed that *T. taiwanensis* and *T. sichuanensis* are indeed distinct species.

**Figure 2.**
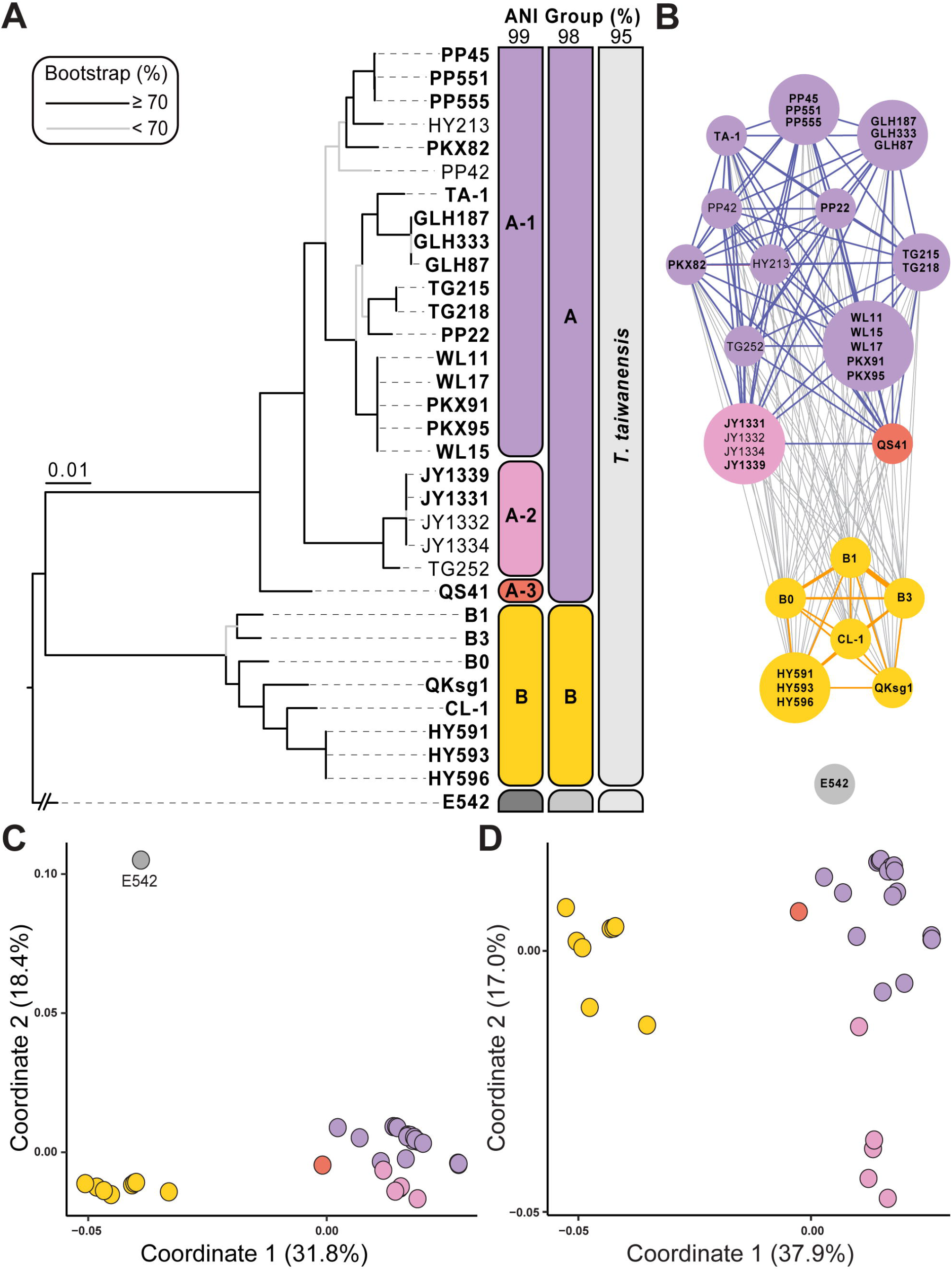
Genomic divergence among the strains analyzed. (A) A maximum-likelihood phylogeny based on 2,159 single-copy genes shared among all strains. The concatenated alignment contains 2,010,937 aligned nucleotide sites. The bootstrap support levels were estimated based on 1,000 re-sampling. Strain E542 from China is included as the outgroup. Strains with complete genome assemblies are highlighted in bold. The grouping of strains based on different cutoff values of genome-wide average nucleotide identity (ANI) are indicated on the right. (B) Population clustering based on gene flow analysis. Strains are consolidated into a single node if their nucleotide sequence divergence is lower than 0.0355%. For each node, the size is proportional to the number of strains and the color is based on 99% ANI grouping. The thickness of each edge reflects the level of gene flow. (C) and (D): Gene content dissimilarity based on principal coordinate analysis, with and without the outgroup, respectively. Each dot represents a strain and the distance between dots reflects the level of gene content differentiation. The % variance explained by each axis is provided in parentheses. The strains are color-coded based on 99% ANI grouping.

In the within-species analysis of these 32 *T. taiwanensis* strains, two distinct populations (i.e., A and B) were identified based on 98% ANI and the core gene phylogeny (**Figure 2A**). When the ANI threshold was increased to 99%, population A could be further divided into three sub-populations: A-1, A-2, and A-3 (**Figure 2A**). Gene flow inference using PopCOGenT (Arevalo et al. 2019) also clustered the 32 *T. taiwanensis* strains into two distinct populations, consistent with the clustering based on 98% ANI (**Figure 2B**). Additionally, no detectable gene flow was found between the outgroup *T. sichuanensis* E542 and any of the 32 *T. taiwanensis* strains. In terms of overall gene content divergence, the two populations of *T. taiwanensis* showed clear differentiation from each other, as well as from the outgroup E542 (**Figure 2C**). When the outgroup was excluded from the analysis to achieve higher resolution in within-species comparisons, all five JY strains belonging to subpopulation A-2 exhibited greater gene content differentiation from other population A strains (**Figure 2D**).

To investigate whether the divergence among these strains is related to isolation by distance, we examined the correlation between geographical distance and two measurements of genomic differentiation, namely the alignment fraction (i.e., the proportion of genomic fragments mapped between two genomes in pairwise comparions) and the ANI values of those mapped fragments. For both measurements, statistically significant results were found in the within-population comparisons (**Figure 3**). Notably, for population B, the geographical distances and ANI values exhibited a strong correlation (R^2^ = 0.52, P = 1.62e^-05^). These results are consistent with a previous study that examined hot spring cyanobacteria divergence at global and local scales (Papke et al. 2003), providing further support that the habitat restriction to hot springs may have shaped the divergence patterns of these bacteria similarly to those observed among island living organisms, rather than the unrestricted disperal expected for free-living bacteria. However, in contrast to the patterns observed for within-population comparisons, the between-population comparisons had only weak (i.e., distance versus alignment fraction; *R*^2^ = 0.05, P = 0.001) or non-significant (i.e., distance versus ANI; *R*^2^ = 0.01, *P* = 0.34) correlations. These findings indicate that while isolation by distance may explain the divergence at within-population level, divergence at higher levels (e.g., between population or species) may be influenced by other factors.

**Figure 3.**
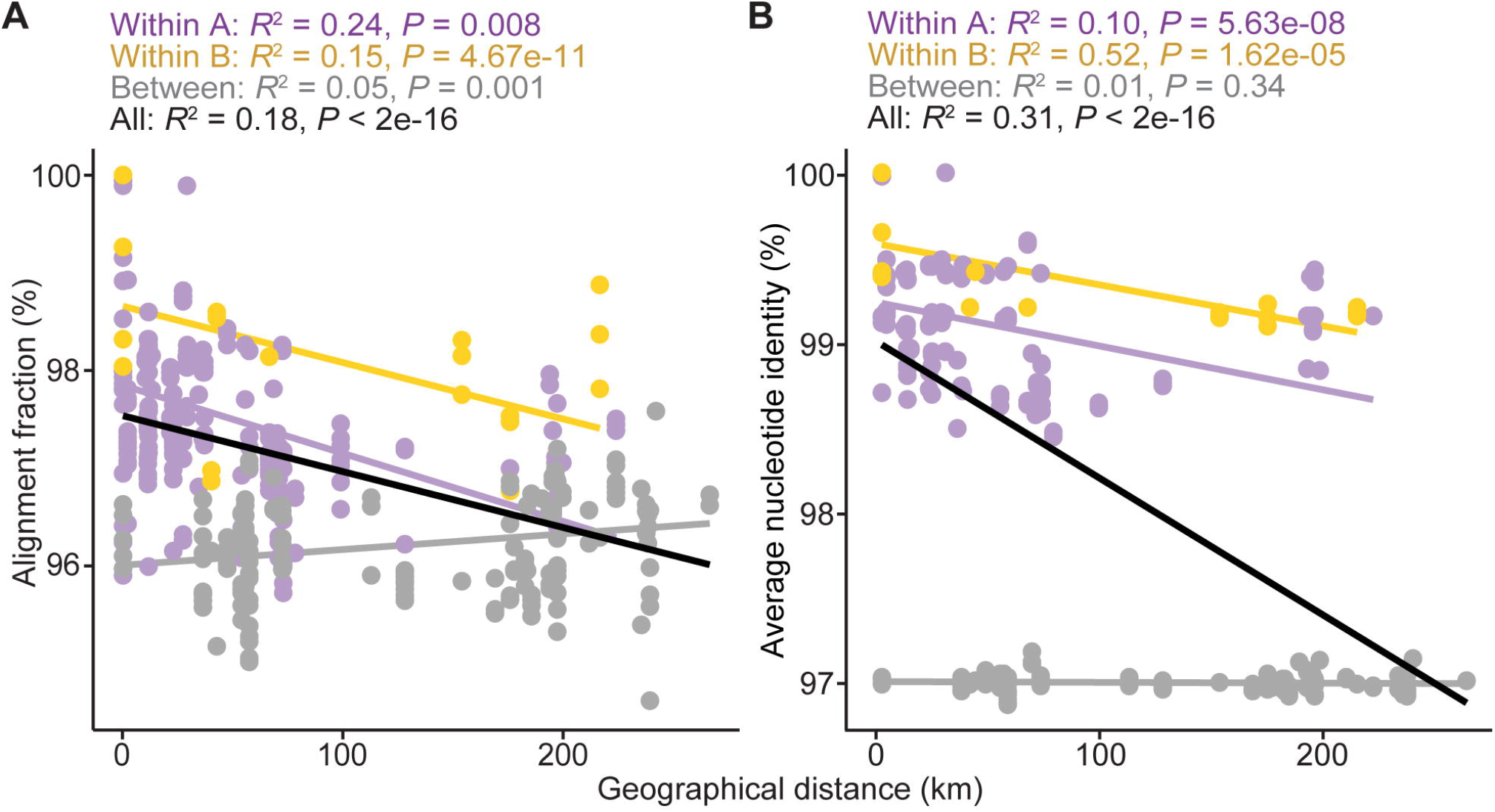
Correlations between geographical distance and genome similarity among the 32 *Thermosynechococcus taiwanensis* strains. (A) Genome similarity measured in alignment fraction (i.e., percentage of genomic fragments mapped in pairwsie comparisons). (B) Genome similarity measured in the average nucleotide identity of the mapped fragments. The data points and linear regression lines are color-coded based on the type of pairwise comparisons: purple, within population A; yellow, within population B; grey, between populations A and B; black: regression line of all data points. The coefficients of determination (*R*^2^) and P values of all linear regression analyses are labeled above the plots.

The pan-genome analysis among these 32 *T. taiwanensis* strains produced a curve of cumulative size that nearly plateaued at 3,030 genes (**Figure 4**). Extrapolation of the sampling curve predicted that adding an additional strain would only increase the pan-genome size by approximately 4.7 genes. Based on Heap’s law (Tettelin et al. 2008), the model fitting parameter γ was estimated to be approximately 0.050. In comparison, recent studies on other cyanobacteria have reported γ values higher than 0.6 (Beck et al. 2018; Cao et al. 2022; Qian et al. 2023). However, those studies were conducted at between-species level and may not be compared directly. For within-species analyses of other bacteria, the estimated value of γ was 0.24 for the environmental bacterium *Leptospirillum ferriphilum* (Zhang et al. 2018) and between 0.12 to 0.5 for pathogenic bacteria (Park et al. 2019; Hyun et al. 2022). These results suggest that our sampling of *T. taiwanensis* strains sufficiently covers for the genomic diversity of this species, and this species likely possesses a closed pan-genome.

**Figure 4.**
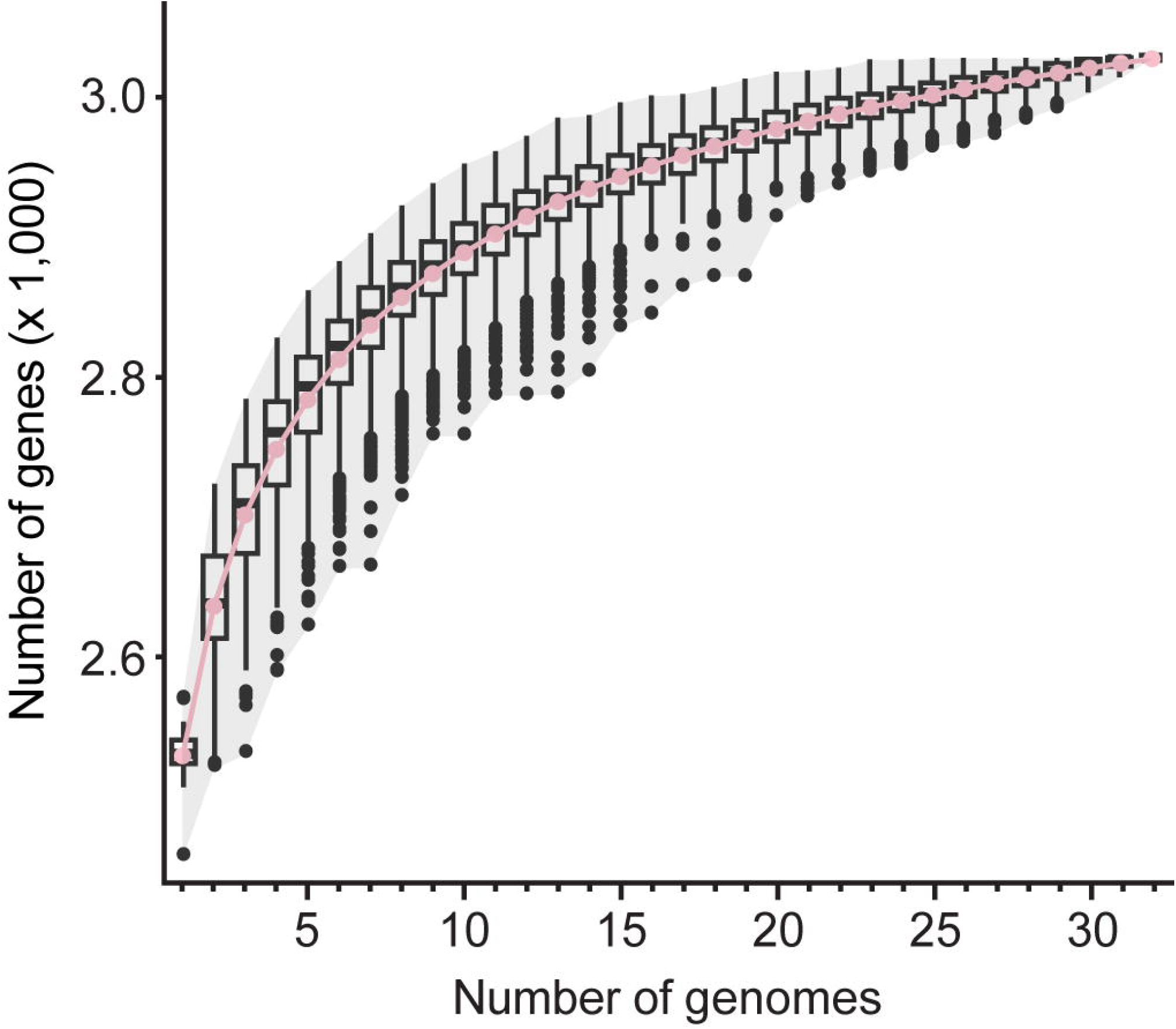
Inference of pan-genome among the 32 *Thermosynechococcus taiwanensis* strains. The box plots illustrate the distribution of 1,000 re-sampling, the pink curve indicates the mean values, the shaded area indicates the range of distribution.

### Inference of selective sweeps

To investigate the evolutionary processes driving within-species divergence of *T. taiwanensis*, we utilized the PopCOGenT (Arevalo et al. 2019) pipeline to infer selective sweeps. The whole-genome alignments were cut into 109,608 sliding windows for maximum-likelihood inference. Regions that passed the filtering steps were merged into 190 loci in population A and 236 loci in population B (**Figure 5A**). Of these loci, 47 in population A and 96 in population B exhibited significantly lower nucleotide diversity (**Figure 5B**), indicating potential selective sweeps following the divergence of these two populations. In total, these loci contained 149 and 289 genes in populations A and B, respectively (**Table S3**). Comparison of the two lists revealed only 16 genes common to both populations (**Table S3**), suggesting that selective sweeps mostly targeted different genes in each population after divergence. Based on the COG classifications, approximately two-thirds of these genes were assigned to specific functional categories (**Figure 5C**). For KEGG analysis, these two sets of genes were assigned to 23 and 29 functional groups, respectively (**Table S3**).

**Figure 5.**
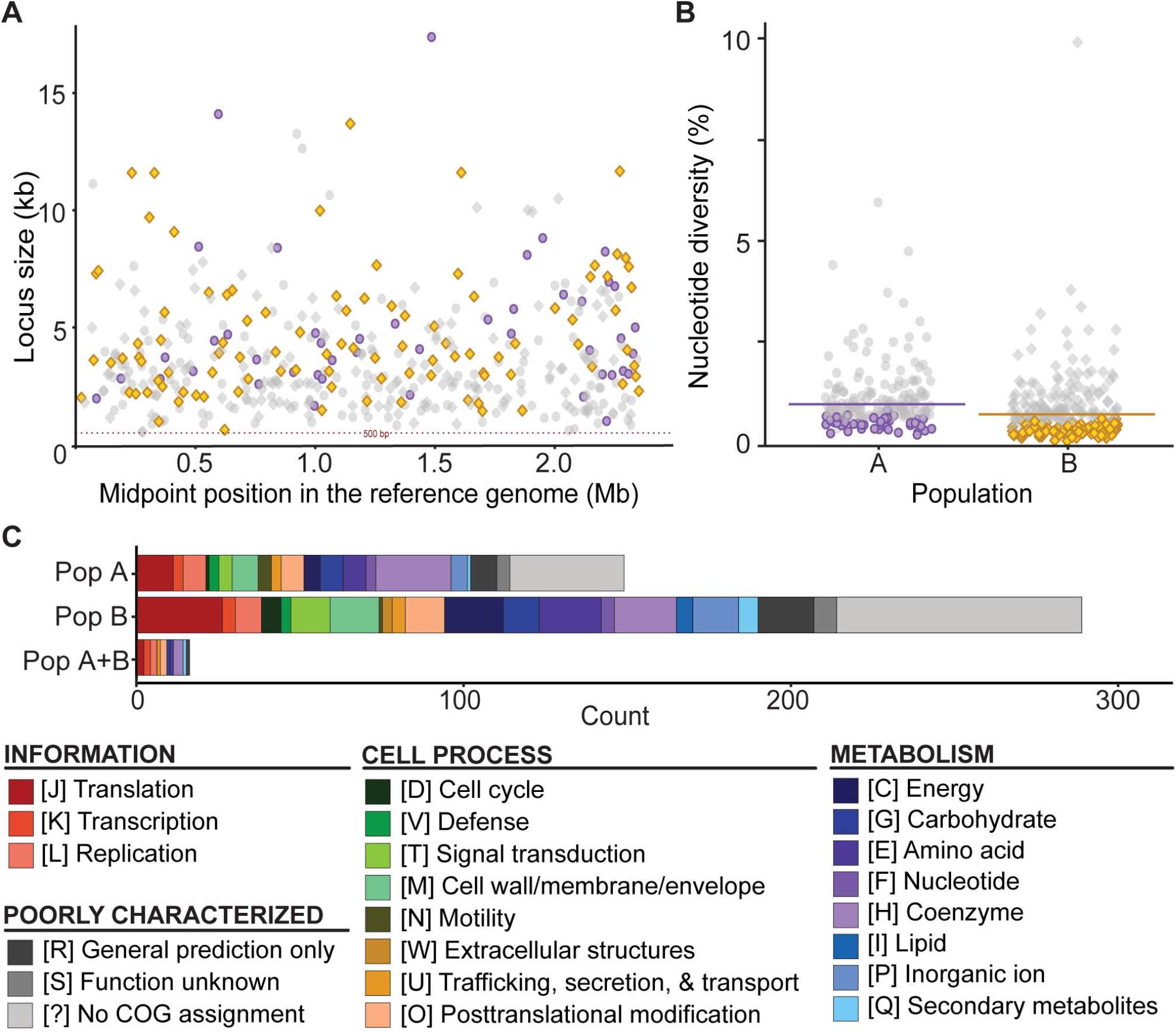
Inference of selective sweeps in the two populations. (A) Sizes and midpoint positions of the loci examined. The genomic positions are plotted based on strain PP45 as the reference. The loci identified as having experienced selective sweeps in populations A and B are illustrated using purple circles and yellow diamonds, respectively. (B) Distribution of nucleotide diversity of all loci in each population. For each population, the horizontal line indicates genome-wide average. The loci identified as having experienced selective sweeps in populations A and B are illustrated using purple circlesyellow diamonds, respectively. (C) Functional classification of the genes identified as having experienced selective sweeps based on COG categories.

Among those genes that were inferred to have experienced selective sweep and can be assigned to specific KEGG functional groups, we highlighted the ones that may be important for ecological adaptation (**Figure 6**). For genes related to photosynthesis (**Figure 6A**), *chlG* (QYC27_02810) that encodes a chlorophyll synthase for the terminal step of chlorophyll biosynthesis was identified as experienced selective sweep in both populations. For genes that experienced selective sweep in only one of the populations, those encode components of allophycocyanin and photosystem II appeared to be major targets. For genes related to motility (**Figure 6B**), *fimT* (QYC27_11160) involved in type IV fimbrial biogenesis, as well as *pilN* (QYC27_11110) and *pilO* (QYC27_11105) involved in type IV pilus assemlby, experienced selective sweep among population A strains. In comparison, genes inferred to have experienced selective sweep among population B strains include those related to two componet systems (*pixH* and *pilG*; QYC27_00350 and QYC27_00355) and type I pilus assembly (*cupE*; QYC27_11185). For genes related to other important functions such as ion transport (**Figure 6C**), exopolysaccharides biosynthesis and transport (**Figure 6D**), fatty acid biosynthesis (**Figure 6E**), and defense systems (**Figure 6F**), only dam (QYC27_02605) that encodes a DNA-methyltransferase for protecting DNA from restriction endonuclease cleavage in type II restriction-modification system was a common target of selective sweep in both populations. For other genes in these functional groups, the targets of selective sweep are more diverse between the two populations. Notably, genes that encode both components of lipopolysaccharide export system, namely *lptA* (QYC27_02640) and *lptB* (QYC27_02645), experienced selective sweep in population A (**Figure 6D**).

**Figure 6.**
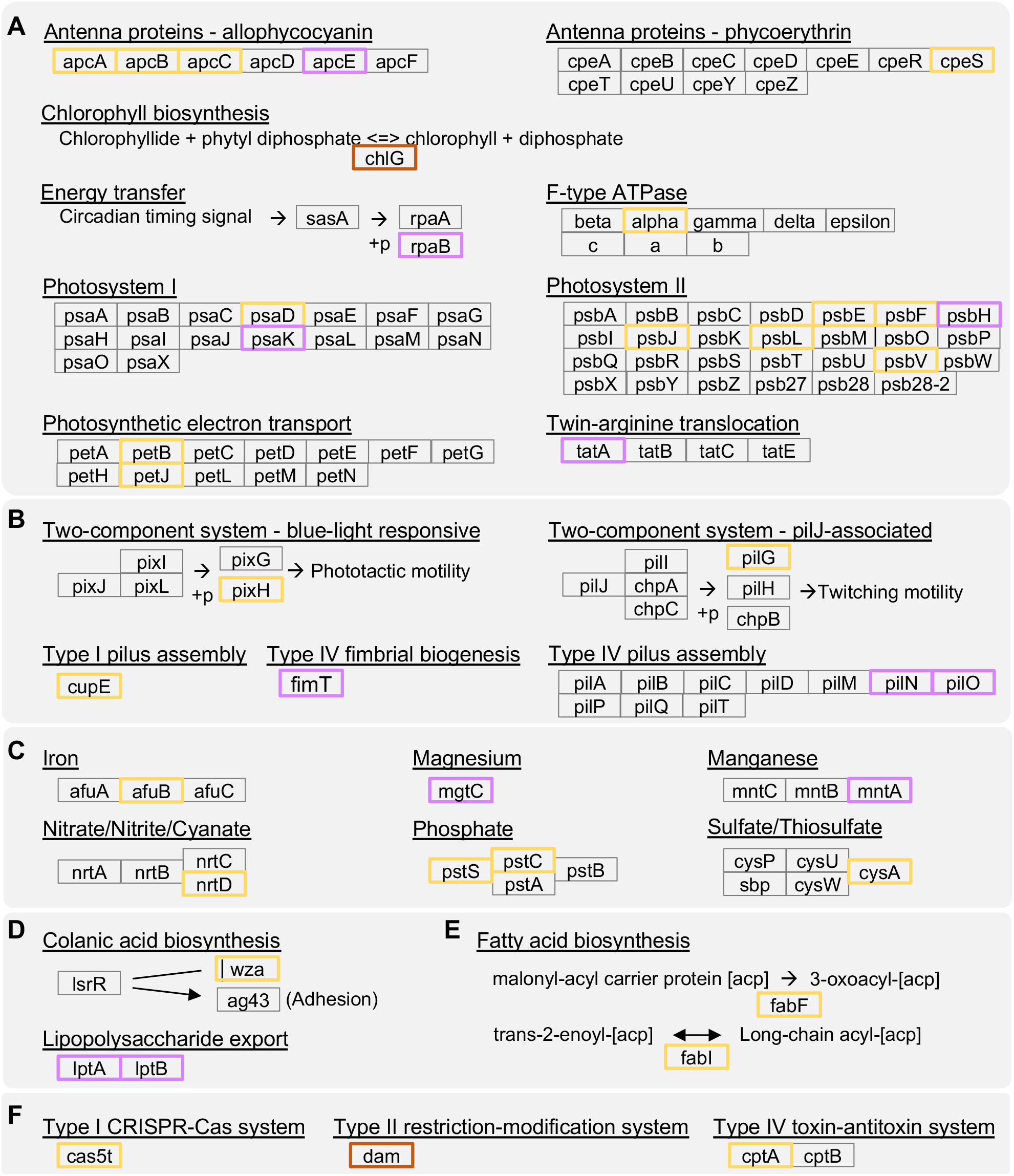
Highlighted examples of genes that experienced selective sweep. The functional groups are based on KEGG pathway classification. The genes that were inferred as experienced selective sweeps in population A, population B, and both populations are highlighted in purple, yellow, and red, respectively. (A) photosynthesis, (B) motility, (C) mineral and organic ion transporters, (D) exopolysaccharides biosynthesis and transport, (E) fatty acid biosynthesis, and (F) defense systems.

Taken together, the inference of these genes provides a list of candidates for future studies to examine the phenotypic divergence between those two populations of strains. If phenotypic divergence is confirmed, these genes are promising targets for comparative functional studies to better understand the ecology and evolution of these bacteria.

## Conclusions

In this work, we conducted extensive sampling of hot springs in Taiwan and found that one species of thermophilic cyanobacterium, namely *T. taiwanensis*, is widely distributed. Moreover, this species likely has a closed pan-genome, and our sampling captured most of its genomic diversity. Analyses based on core-genome phylogeny, gene flow estimates, and overall gene content divergence all supported the within-species divergence into two major populations. While the specific environmental factors associated with the population divergence remain unclear, the inference of genes that have experienced selective sweep offers valuable clues for future studies to better understand the genetic basis of intra-specific divergence and potential speciation. In addition to advancing microbial genomics and evolution, the strains collected and the genome sequences produced in this study provided valuable resources for future research.

## Declarations

## Ethics approval and consent to participate

Not applicable.

## Consent for publication

Not applicable.

## Availability of data and materials

The genome sequences reported in this work have been deposited in NCBI GenBank under the accession numbers GCA_030518695.1, GCA_030518715.1, GCA_030518735.1, GCA_031432345.1, GCA_031460315.1, GCA_031460915.1, GCA_031460935.1, GCA_031460975.1, GCA_031583185.1, GCA_031583205.1, GCA_031583225.1, GCA_031583255.1, GCA_031583295.1, GCA_031583315.1, GCA_031583335.1, GCA_031583355.1, GCA_031583375.1, GCA_031583395.1, GCA_031583415.1, GCA_031583435.1, GCA_031583455.1, GCA_031583475.1, GCA_031583495.1, GCA_031583515.1, GCA_031583535.1, GCA_031583555.1, GCA_031586415.1. More detailed information is provided in **Table S2**. One representative strain, JY1331, has been deposited in the Bioresource Collection and Research Center (BCRC) in Taiwan under the accession number 81421.

## Authors’ contributions

HYC collected the samples, conducted the experiments and analysis, prepared the figures and tables, and wrote the draft. HCY validated the analysis and contributed to the preparation of the draft. HAC helped with the initial planning, provided technical assistance, and revised the draft. CHK conceptualized and supervised the project, acquired the funding, and wrote the draft. All authors read and approved the final manuscript.

## Acknowledgements

The Sanger sequencing, whole-genome shotgun sequencing library preparation, and Oxford Nanopore Technologies sequencing services were provided by the Genomic Technology Core (Institute of Plant and Microbial Biology, Academia Sinica, Taiwan). The Illumina NovaSeq sequencing service was provided by Genomics BioSci and Tech Co, Ltd (New Taipei, Taiwan). We thank Shu-Jen Chou, Ai-Ping Chen, Mei-Jane Fang, Ming-Ling Cheng, members of the Plant and Environmental Microbiology Group in our institute, and members of the Academia Sinica Evolution Club for technical assistance and scientific discussions.

## Supplementary Information

Additional file 1: **Table S1**. List of the sampling sites.

Additional file 2: **Table S2**. List of the genome assemblies.

Additional file 3: **Table S3**. Lists of the genes inferred as experienced selective sweeps. The locus tags are based on strain PP45.

